# Flexible estimation of biodiversity with short-range multispectral imaging in a temperate grassland

**DOI:** 10.1101/2022.03.08.483493

**Authors:** J. Jackson, C. S. Lawson, C. Adelmant, E. Huhtala, P. Fernandes, R. Hodgson, H. King, L. Williamson, K. Maseyk, N. Hawes, A. Hector, R Salguero-Gómez

**Affiliations:** Department of Biology, University of Oxford, UK, OX1 3SZ; School of Environment, Earth & Ecosystem Sciences, The Open University, UK, MK7 6AA; Oxford Robotics Institute, Department of Engineering Science, University of Oxford, UK, OX1 3PJ; Max Planck Institute for Demographic Research, Rostock, Germany, DE 18057

**Keywords:** Autonomous monitoring, Biodiversity Drone, Remote sensing, Unmanned Aerial Vehicle (UAV)

## Abstract

1. Image sensing technologies are rapidly increasing the cost-effectiveness of biodiversity monitoring efforts. Species differences in the reflectance of electromagnetic radiation have recently been highlighted as a promising target to estimate plant biodiversity using multispectral image data.
2. However, these efforts are currently hampered by logistical difficulties in broad-scale implementation and their use in characterizing biodiversity at different spatial scales.
3. Here, we investigate the utility of multispectral imaging technology from commercially available unmanned aerial vehicles (UAVs, or drones) in estimating biodiversity metrics at short-range (<10 m image recording height) in a temperate calcareous grassland ecosystem in Oxfordshire, UK. We calculate a suite of moments (coefficient of variation, standard deviation, skew, kurtosis) for the distribution of radiance from multispectral images at five wavelength bands (Blue 450±16 nm; Green 560±16 nm; Red 650±16 nm; Red Edge 730±16 nm; Near Infrared 840±16 nm) and test their effectiveness at estimating ground-truthed biodiversity metrics from *in-situ* botanical surveys for 37 - 1 m × 1 m quadrats.
4. We find positive associations between the average coefficient of variation in spectral radiance and both the Shannon-Weiner and Simpsons biodiversity indices. Furthermore, we find that the average coefficient of variation in spectral radiance is consistent and highly repeatable, across sampling days and recording heights. Positive associations with biodiversity indices hold irrespective of the image recording height (2-8 m), but we report reductions in estimates of spectral diversity with increases to UAV recording height. UAV imaging reduced sampling time by 16-fold relative to *in-situ* botanical surveys.
5. *Synthesis* - We demonstrate the utility of multispectral radiance moments as an indicator of grassland biodiversity metrics at high spatial resolution using a widely available UAV monitoring system at a coarse spectral resolution. The use of UAV technology with multispectral sensors has far-reaching potential to provide cost-effective and high-resolution monitoring of biodiversity in complex environments.

## 1. Introduction

With over 1 million species expected to go extinct by 2100, cost-effectively monitoring biodiversity is a critical task in the Anthropocene (Díaz et al., 2019). Image sensing technologies, which can be used to monitor biological systems through the measurement of reflected and emitted radiation, have emerged as a critical tool that can increase this cost-effectiveness (Turner, 2014). The characterisation of floral biodiversity with remote sensing, particularly with satellite imagery, is well-established in biodiversity research (Pettorelli et al., 2005). Multiple efforts have been made towards using remote sensing data, particularly at large spatial scales and in forest ecosystems, to estimate plant diversity (e.g., Frye et al., 2021; Jetz et al., 2016; Tuanmu & Jetz, 2015; Turner et al., 2003). However, there are limitations in the use of long-range remote sensing, including low spatial resolution that does not necessarily highlight biodiversity at small spatial scales (Mairota et al., 2015), high sensor costs (*e.g*., $98,700, Headwall Photonics, 2022) and monitoring costs (*e.g*., $60,000, Jet Propulsion Laboratory, 2022), and reliance on publicly available satellite data (e.g., The European Space Agency, 2022). Flexible application of remote sensing concepts and technology at a wide range of spatial scales, in complex changing environments, and with increased cost-effectiveness, will provide vital resources for monitoring biodiversity (Turner, 2014).

Reflectance of electromagnetic (EM) radiation beyond the visible range (380-700 nm) has recently been shown accurate proxy for biodiversity (Cavender-Bares et al., 2020; Wang & Gamon, 2019). The general concept of ‘spectral diversity’ is founded on the principle that, due to differences in functional form (both growth form and pigmentation), plant species have differential reflectance signals across the electromagnetic (EM) spectrum (Gamon et al., 1997). Thus, for a multispectral image, the diversity of spectral reflectance can be a proxy for the number of different plant species, or species diversity (Gholizadeh et al., 2019; Laliberté et al., 2020). The spectral diversity concept was recently applied in the hyper-diverse Cape Floristic Region, where destructively sampled leaf reflectance spectra were used to obtain a robust proxy (*R*^2^ > 0.9) of species diversity across 1,267 - 10 m × 5 m quadrats (Frye et al., 2021). Therefore, integrating sensing data at a range of spatial scales (Laliberté et al., 2020; Turner, 2014) and the use of spectral surrogates for biodiversity (Frye et al., 2021) can rapidly improve biodiversity monitoring.

Recently, there have been several applications of spectral diversity from close range imaging data (Gholizadeh et al., 2019; Lopatin et al., 2017). In prairie grassland ecosystems, close associations have been found between species diversity and spectral diversity, captured using aircraft-mounted hyperspectral sensors and images at a spatial resolution of 1 m × 1 m (pixel resolution) (Gholizadeh et al., 2018, 2019, 2020). Gholizadeh et al. (2019) primarily use the average coefficient of variation across pixels and spectral bands as the metric of spectral diversity, which we also adopt here. Furthermore, at a fine resolution of < 0.5 cm × 0.5 cm, static monitoring of grassland plots has been used to estimate not only biodiversity metrics (Imran et al., 2021; Villoslada et al., 2020), but to reconstruct species percentage cover and extract detailed features of community dynamics (Lopatin et al., 2017). However, a key limitation of these close-range imaging approaches is their reliance on expensive hyperspectral sensors (> $50,000 sensors; Gholizadeh et al., 2019; Imran et al., 2021; Lopatin et al., 2017) and monitoring ($1,200 per hour using the CALMIT aerial sensor from Gholizadeh et al., 2019). Overcoming these cost limitations will facilitate further use of spectral imaging in biodiversity research.

Despite advances in image sensing, there is a need for monitoring systems that i) are deployable in a diverse range of habitats, ii) can span a range of spatial resolutions, iii) are cost-effective, and iv) user friendly. One potential solution is the use of commercial unmanned aerial vehicles (UAVs), or drones, which have rapidly increased in popularity over the last decade (Colomina & Molina, 2014). Here, we investigate the efficacy of multispectral imaging UAV technology in the estimation of biodiversity in a temperate calcareous grassland. We used a commercially available UAV system with a five-band multispectral band sensor (Blue 450±16 nm; Green 560±16 nm; Red 650±16 nm; Red Edge 730±16 nm; Near Infrared 840±16 nm) to image 37 - 1 m × 1 m quadrats that were also characterised using *in situ* biodiversity assessments from botanical surveys. Then, by extracting moments of spectral diversity from multispectral image data, we estimate spectral diversity and explored associations between spectral diversity and biodiversity. Our goal is to provide a framework for cost-effective biodiversity monitoring in complex environments using readily and commercially available technologies.

## 2. Materials and Methods

### 2.1 Study site and in-situ biodiversity data

Data collection took place at the five-acre section of the Upper Seeds field site (51°46’16.8”N 1°19’59.1”W) in Wytham woods, Oxfordshire, UK between 16^th^ June-14^th^ July 2021, the peak of the growing season. The Upper Seeds site is a recovering and managed calcareous grassland, which was used for agriculture in the 1950s, before encroaching scrub vegetation was removed and the site was managed as a grassland beginning in 1978 (Gibson, 1986). Management on upper seeds is implemented with mowing of the site in mid-July at the peak of the growing season, and again in early October, coinciding with the end of the growing season. The site has a low average soil depth (300-500 mm), generally alkaline soils (Gibson & Brown, 1991), a daily average temperature range of −5 °C to 26 °C (2016-2020), a daily total precipitation range of 0-40 mm (2016-2020), and high general biodiversity, in which graminoids are the dominant functional group (59.1% by biomass). A total of 37 1 m × 1 m experimental quadrats were used in the current study, which display a large degree of variation in species composition and biomass. There were between 16 and 33 vascular plant species per m^2^, with a mean richness of 25.77 species and a median richness of 26 species. Total above-ground dry biomass across quadrats varied between 166.8 g/m^2^ and 931.5 g/m^2^, with a mean of 397.9 g/m^2^ and median of 327.2 g/m^2^.

We explored biodiversity and spectral diversity associations in the context of two long-term experiments that aim to explore the response of grasslands to environmental change (full site map in Fig. S1). These experiments are the Disturbance and Resources Across Global Grasslands (DRAGNet, n = 20 plots) coordinated research network (https://nutnet.org/dragnet) and the global drought network (DroughtNet, n =17 plots) coordinated research network (https://drought-net.colostate.edu/). All DRAGNet plots (5 m × 5 m plots) were ambient controls, with no experimental treatments applied prior to the collection of the data reported here. Each 5 m × 5 m plot from DroughtNet was one of four experimental treatments: ambient control plots (n = 5), −50% rainfall shelters to simulate drought (n = 5), +50% irrigated plots to simulate increased rainfall (n = 5), and procedural controls (rainfall shelter with no change to rainfall) (n = 2; three plots were inaccessible for the UAV as the rainfall shelters were fixed). For analyses, ambient control treatments (n = 25) across the two research networks were pooled as we did not observe substantial differences between biodiversity metrics (Fig. S2). To account for replicated observations of the same quadrats and estimate the consistency of spectral diversity measures, we explored quadrat-level variance using intercept-only random effects for the quadrat ID.

To estimate the efficacy of multispectral sensors in predicting biodiversity, we collected data from two sources, *in-situ* biodiversity assessments and UAV derived multi-spectral image data (Fig. 1). For the *in-situ* assessments, we quantified biodiversity metrics using species-level percentage cover and dry above-ground biomass data. We estimated percentage cover data for all vascular plant species in a plot using a 1 m × 1 m gridded quadrat (10 cm grid), focusing on four broad functional groups: graminoids, legumes, forbs, and woody species (Fig. 1). Because species overlapped spatially, percentage cover estimates could exceed 100%. Using relative proportions, *p*, calculated from percentage cover estimates, we calculated three biodiversity metrics: i) vascular plant species richness, ii) the Shannon-Weiner diversity index, *H* (equation 1; Shannon & Weaver, 1963), and iii) the Simpson’s diversity index, *D* (equation 2; Simpson, 1949):

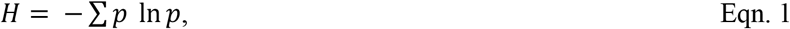

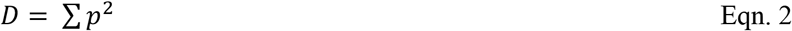

**Fig. 1.**
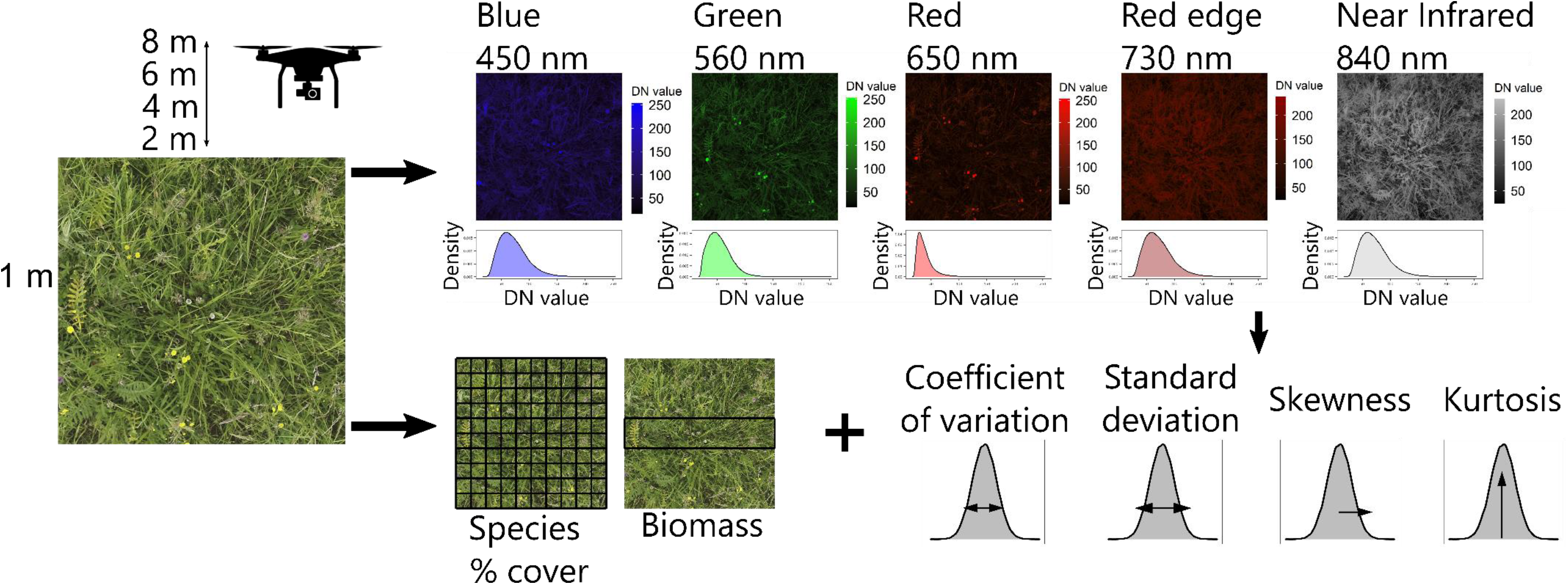
Schematic for assessing the efficacy of spectral distribution moments for capturing biodiversity in a temperate grassland. For each 1m × 1 m observation quadrat, we collected both UAV image data (top) and in-situ biodiversity data (bottom). In-situ biodiversity data were collected by botanical surveys for vascular plant percentage cover across the quadrat (from which richness, Shannon-Weiner and Simpson’s indices were calculated) and using dry above-ground biomass for clip strips (area determined by the coordinated research networks DRAGNet and DroughtNet; see 2.1), after UAV images were taken. UAV images were collected for each plot at four recording heights (2 m, 4 m, 6 m, and 8 m) across five multispectral bands for which at-sensor radiance Digital Number (DN) value distributions were summarised using four moments. Finally, *in situ* biodiversity data were combined with spectral distribution moments to examine their potential relationships using a Bayesian linear regression framework.

We estimated above ground biomass after UAV image/percentage cover data collection, using a clip strip of all vascular plant material in 1 m × 0.2 m (DRAGNet; collected from standardised locations in the plot) or 1 m × 0.25 m (DroughtNet; collected from the centre of each quadrat). Clip strips were gathered using hand trimmers at a height of 1-2cm above the soil surface. Within one day of collection, we sorted clip strips in to five functional groups: graminoids, legumes, forbs, woody species, and bryophytes (not included in species-level percentage cover estimates) and dried them at 70°C for 48hr, before weighing the dry biomass at an accuracy of ±0.1g. The estimates of biomass were scaled to g/m^2^ for analyses.

### 2.2 UAV image data collection

To obtain spectral diversity data, we collected image data using manual flights of the DJI Phantom 4 multispectral UAV (https://www.dji.com/p4-multispectral). The sensor payload of the DJI Phantom 4 multispectral consists of six 4.96 mm × 3.72 mm complementary metal–oxide–semiconductor (CMOS) sensors: one RGB sensor for visible range colour images, and five monochrome sensors for multispectral imaging. The five multispectral sensors are sensitive at the following electromagnetic wavelengths: Blue - 450 nm ± 16 nm, Green - 560 nm ± 16 nm, Red - 650 nm ± 16 nm, Red edge - 730 nm ± 16 nm and Near-infrared - 840 nm ± 26 nm (Fig. 1). Each sensor has an effective resolution of 2.08 MP. All six image sensors are triggered simultaneously when capturing data, with negligible (but non-0) positional differences between sensors. A dorsal spectral sunlight sensor on the P4 multispectral sensor provides image exposure compensation of multispectral image data.

To obtain spectral diversity metrics, we collected multispectral images for each quadrat over several flights across the sampling period, capturing quadrat-level variability with weather/light conditions. However, to minimise visual interference (from rain or low sun), all images were taken during dry weather and between 10:30-15:30 (BST). The corners of each DRAGNet quadrat were marked with flags (Fig. S1b). For DroughtNet, the quadrat was approximated using the outer edges of the 5 m × 5 m plot. All images were collected facing the western edge of each plot. We collected images at increasing approximate image recording heights of 2 m, 4 m, 6 m, and 8 m above the ground to capture changes in image resolution and consequences for estimating biodiversity (Fig. 1). Flying height is recorded relative to the UAV’s take-off location, and although the topographical variation at the site is < 5 m, image record heights were approximated using structures of known height (*i.e*., rainfall shelters, see Fig. S5). A total of 1,878 individual images were collected for the 37 quadrats over seven sampling days.

### 2.3 Image processing

To extract spectral reflectance metrics from the raw image data, we standardised raw images across quadrats for each sample. The raw images encompassed the full field of view of the sensors, and we first batch-cropped images with Adobe Lightroom v. 5 (Adobe, 2021) to include only data for the desired 1 m^2^ quadrats. Images were exported as .tif files maintaining at-sensor radiance values with minimal post-processing. We did not perform post-processing to account for interference from other sources solar radiation (e.g. Gholizadeh et al., 2019) due to the low recording height of UAV flights in the current study. Instead we estimated quadrat-level variability in spectral signals across the study period. The data stored in each .tif image were at-sensor radiance Digital Number (DN) values, which were used in subsequent analyses. The use of raw DN values was appropriate for the purpose of this study because we aimed to capture moments of the distribution of DN values in each image. Scaling DN values to normalised reflectance values was not necessary to explore the radiance distributions because relative differences between DN values, and thus distributional moments remain unaltered. Furthermore, the omission of calibration to reflectance values, which is typically performed with standardised reflectance plates, was more appropriate for the application of the current protocol and aims *i.e*. deployment in complex environments. Multispectral .tif images were treated as rasters for further image processing, and all subsequent analysis was carried out using R version 4.0.5 (R Core Team, 2021).

We calculated moments of spectral radiance for each image using the *raster* package (Hijmans, 2020). Following Gholizadeh et al. (2019), we calculated the coefficient of variation, standard deviation, skewness, and kurtosis across raster pixels to capture the shape of the at-sensor radiance DN distribution (Fig. 1). We averaged moment values of radiance DNs across all multispectral bands for a single observation (a given quadrat at a given recording height in each sampling event) to calculate overall distributional moments. Thus, here we define the spectral coefficient of variation as the mean coefficient of variation of the spectral radiance across raster pixels and multispectral bands for a single image. Observations were discarded if the image recording height was >8 m and replicate images were not obtained for all quadrats at all image recording heights. Therefore, the final sample size for the averaged spectral moment data was 193. In addition, to identify the spectral bands that were most sensitive to biodiversity metrics, we also tested band-level associations, where radiance distributions were not averaged across spectral bands for the same raw data, and in this case spectral radiance distributions of each band were related to biodiversity indices.

### 2.4 Statistical analyses

We explored the efficacy of spectral radiance distribution moments in describing *in-situ* biodiversity indices using a Bayesian hierarchical linear regression model selection framework using the *brms* package (Bürkner, 2017; Fig. 1). All variables were z-scored (mean and variance centered on 0) for analysis to meet the distributional assumptions of linear regressions. The key response variable was the spectral coefficient of variation (Gholizadeh et al., 2019), and the key predictor variables were the *in-situ* biodiversity indices. However, we also tested other spectral moment-biodiversity associations, namely, the skewness of spectral radiance and biomass.

We then estimated the out-of-sample predictive performance of models including biodiversity indices relative to base models. For each explored pair-wise combination of spectral distribution moment and biodiversity indices, we performed leave-one-out cross validation with the *loo* criterion and the expected log-wise predictive density (*elpd*, where Δ*elpd* gives the change in *elpd* relative to another explanatory model) (Vehtari et al., 2017). Base models did not include any predictor variables, including only an intercept-only random effect for quadrat. We also investigated the performance of models including image recording height, and two-way interaction terms between biodiversity indices and height to explore how image resolution change influenced the efficacy of spectral diversity indices.

In addition to models on averaged spectral moments, we also used band-level moments to investigate the relationship between biodiversity indices and the spectral coefficient of variation for each individual EM band. We included univariate and two-way interaction terms between the EM band and biodiversity indices variables. Finally, because a small number of quadrats used in the current study were also exposed to long-term drought/irrigation/control treatments, we explored whether there were differences in average spectral moments between ambient (n = 25 plots), control (n = 2), irrigated (n = 5) and drought (n = 5) using a categorical predictor for treatment.

To account for repeated observations from the same quadrat at different heights or across sampling events, all models included an intercept-only random effect for the quadrat *σ_quadrat_*. From this random effect, we also estimated the intraclass correlation (ICC) or repeatability (*R*). This term indicates the proportion of quadrat-level variance *σ_quadrat_* with respect to the population-level variance *σ* (Nakagawa & Schielzeth, 2010). We used this estimate of repeatability enabled us to assess the consistency of spectral radiance distributions across observations of the same quadrat.

In analyses with average spectral distribution moments, we used weakly informed normal priors for the population-level intercept and coefficient terms of *N*(0,1). The *σ_quadrat_* term was fit using an exponential prior with a rate of two. For band-level analyses (with a greater number of parameters), models were fit using *N*(0,0.7) intercept/coefficient priors and exponential *σ_quadrat_* priors with a rate of four. Models were run across four serial chains for 2,000 iterations with 1,000 warmup iterations, and the model’s convergence across chains was assessed by inspecting 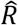 values (Bürkner, 2017).

## 3. Results

We found consistent positive associations between the average coefficient of variation in spectral radiance and biodiversity, namely, the Shannon-Weiner and Simpson’s indices (Fig. 2). The model including the Shannon-Weiner index and image recording height as univariate terms outperformed the base model, with Δ*elpd* = 123.6 (Table S1). Increases in the Shannon-Weiner index were associated with increases in the average spectral coefficient of variation (*β_shannon_* = 0.19 [−0.04, 0.43], 95% credible intervals; Fig. 2a). Furthermore, as expected, there was a strong negative association between image recording height and the average spectral coefficient of variation (*β_height_* = −0.28 [−0.31, −0.26]; Fig. 2a). This negative association suggests that the resolution of spectral diversity decreases rapidly with recording height at this spatial resolution (~ 30 % decrease in scaled spectral variation per 1 m height increase). The positive association with the spectral coefficient of variation was stronger for the Simpson’s biodiversity index (Fig. 2b). Although the full model including a two-way interaction between recording height and Simpson’s index was the best predictive model, we selected the model including only univariate effects (Δ*elpd* = 125.0), because of a lack of a clear interaction effect (Table S2). Here, a similar patten with image recording height was accompanied by a stronger positive association between Simpson’s index and spectral coefficient of variation (*β_simpsons_* = 0.33 [0.12, 0.54]; Fig. 2b).

**Fig. 2.**
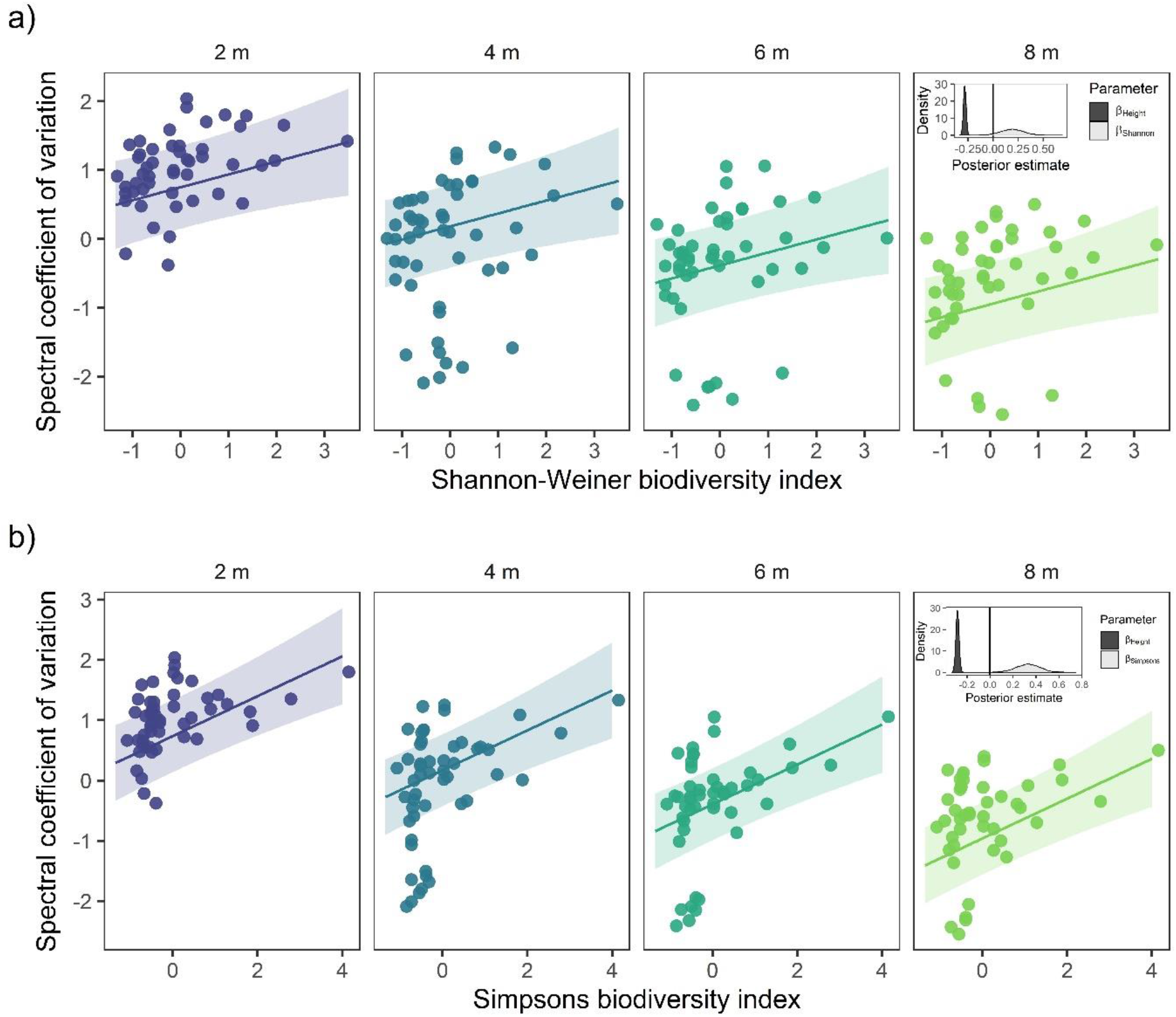
Consistent positive associations between biodiversity indices and the average spectral coefficient of variation. The positive association between (a) Shannon-Weiner biodiversity index and (b) Simpsons index, and the average spectral coefficient of variation (averaged across five spectral bands) for different image recording heights: 2 m, 4 m, 6 m, and 8 m (panels). Points are observations from a single quadrat at a given height. Both biodiversity indices and spectral coefficient of variation are z-scored. Lines represented the posterior prediction mean over 4,000 simulations averaged over all quadrats, with the 90% credible intervals. Insets showcase density distributions of the posterior estimates for the image recording height (*β_height_*) and biodiversity indices (*β_shamon_* and *β_height_*).

Generally, we did not observe associations between the skewness and kurtosis in spectral radiance distributions and biodiversity indices (Fig. S3). Furthermore, there was no clear evidence for a relationship between total above-ground biomass and any of the spectral distribution moments (Fig. S3). Specifically, although there was increased model predictive performance from models including biomass, there was no clear relationship between the skewness of spectral radiance distribution and biomass (*β_biomass_* = 0.08 [−0.19, 0.35]; Fig. S3; Table S5)

In addition to overall effects, in band-level analyses where raw data were not averaged across bands, there was evidence for an interaction effect between the spectral band and both the Shannon-Weiner (Δ*elpd* = 772.0) and Simpson’s indices (Δ*elpd* = 771.5) (Table S3 & S4, respectively). Generally, the green (560±16 nm) and red (650±16 nm) spectral bands displayed higher variability in the coefficient of variation across quadrats, and stronger associations with the Shannon-Weiner and Simpson’s indices (Fig. 3). The Red Edge (730±16 nm) and Near Infrared (840±16 nm) bands exhibited weaker associations with biodiversity indices (Fig. 3).

**Fig. 3.**
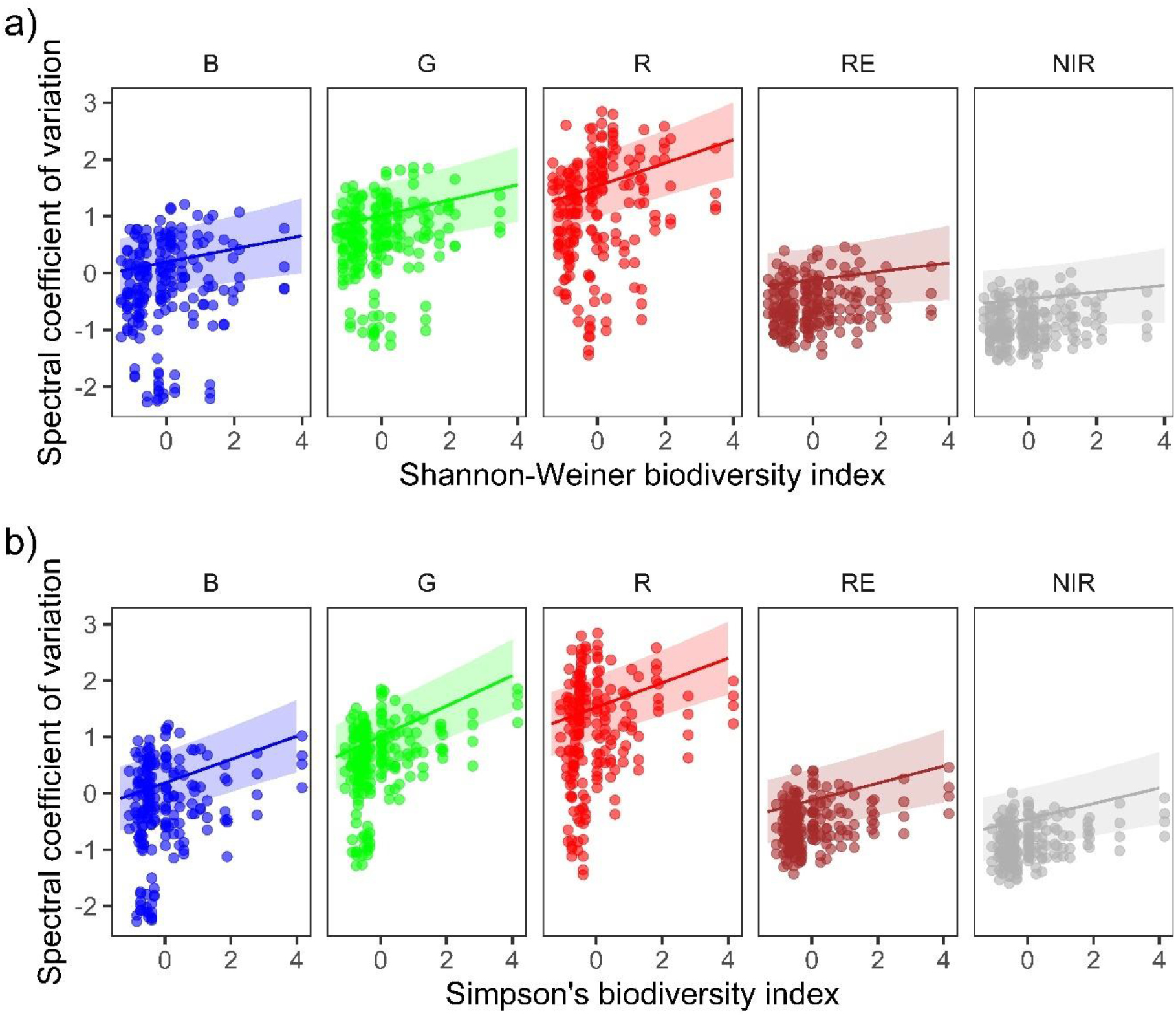
Green and Red spectral bands are the most sensitive to biodiversity indices. Posterior predictions for the band-level spectral coefficient of variation with the Shannon-Weiner (a) and Simpson’s (b) biodiversity indices at an image recording height of 2 m. Here, raw spectral moments were not averaged across spectral bands (as in Fig. 2). Points are raw observations from a single plot, and the colour denotes the spectral band (B = Blue, G = Green, R = Red, RE = Red Edge, NIR = Near Infrared). Both biodiversity indices and spectral coefficient of variation are z-scored. Lines are the posterior prediction mean over 4,000 simulations averaged over plots, with the 90% credible intervals.

When assessing the influence of treatment on spectral radiance, we also observed reductions in the average spectral coefficient of variation in both drought and procedural control quadrats in comparison to ambient or irrigated treatments (Fig. S4). However, given the congruence of procedural control and drought treatments in DroughtNet, both of which are characterised by metal rainfall shelters, we conclude that the reduction in spectral coefficient of variation in drought and procedural treatments is likely a result of structural interference from the rain shelter structures (Fig. S5).

Finally, we tested the consistency of the spectral coefficient of variation across observation days and heights for each quadrat in the best predictive Shannon-Weiner and Simpson’s index models (Table S1 & S2, respectively). The average coefficient of variation was highly consistent for each quadrat when images at different heights or across sampling events were compared (Fig. S6). Both the Shannon-Weiner and Simpson’s models with the average coefficient of variation exhibited quadrat-level variance that exceeded the population-level variance and a high degree of repeatability (Fig. S5; 0.76 [0.65, 0.85] and 0.72 [0.60, 0.82], respectively).

## 4. Discussion

Despite rapid technological advancements in image sensing over the last four decades, biodiversity monitoring is not currently able to track the full extent of human impacts on the biosphere (Wang & Gamon, 2019; Wilson, 2017). We urgently need more cost-effective and widely available systems to monitor detailed changes in biodiversity (Turner, 2014; Turner et al., 2003). Here, using a commercially available short-range Unmanned Aerial Vehicle (UAV, drone), we find a consistent, repeatable association between variation in spectral radiance and species diversity in a biodiverse temperate grassland. The coefficient of variation in spectral radiance was positively associated with the Shannon-Weiner and Simpsons indices, and in particular the green and red bands of the electromagnetic (EM) spectrum were most indicative of grassland biodiversity. Our results build on extensive work in grassland ecosystems exploring the use of spectral diversity as a surrogate for biodiversity (Frye et al., 2021; Gholizadeh et al., 2019; Villoslada et al., 2020) and species composition (Lopatin et al., 2017). However, our research in a diverse temperate grassland community contrast with previous findings that highlighted limitations to the characterisation of biodiversity using spectral imaging in species-rich environments (Imran et al., 2021). We highlight the importance of close-range remote sensing for biodiversity monitoring (Turner, 2014; Turner et al., 2003; Wang & Gamon, 2019). Crucially, we demonstrate the feasibility of the spectral diversity concept using a commercially available UAV with a low spectral resolution sensor, which has far-reaching potential as a tool to explore biodiversity change at high spatio-temporal resolution and in complex environments.

The key advantage of using commercially available UAV technology is its cost-effectiveness relative to reliance on *in-situ* monitoring or the use of long-range or high spectral resolution sensors. In the current study, where we collected data from 37 quadrats at four different image recording heights, the total flight time was 134 min. If we reasonably allocate one researcher 60 min to do a full botanical survey (species percentage cover and biomass clip – ignoring biomass processing of 30 min per sample), the full *in-situ* sampling time is 37 hr. Thus, remote monitoring would have saved on sampling time by a factor of 16. In the current study, the reduction in sampling time is of course limited to proxies of broad biodiversity metrics. However, with increases to sensor resolution and decreasing costs, reconstructing species-level biodiversity data with flexible remote monitoring may also be possible (Lopatin et al., 2017). Increases in cost-effectiveness may also be increased by automated flight paths over survey locations, for which unsupervised spectral reflectance data could be collected. Furthermore, while rapid advancements have been made on spectral diversity, previous studies have utilised high-resolution multi/hyperspectral sensors that are either immobile, high-cost, or both (Frye et al., 2021; Gholizadeh et al., 2019; Imran et al., 2021; Lopatin et al., 2017). The current complete monitoring system is available for purchase for < 10,000 USD, relative to >50,000 USD for many hyperspectral sensors. Moreover, because the most sensitive spectral radiance moments for biodiversity indices were in the green and red bands, which lie within the visible range, we provide evidence that spectral radiance from visible-only imaging may be sufficient to act as a proxy for biodiversity indices. The present method provides a low-cost and mobile solution to rapidly characterise biodiversity in complex environments.

The rapid increase in the public use of drone technology provides an opportunity for a rapid expansion of detailed biodiversity monitoring at flexible spatio-temporal scales (Colomina & Molina, 2014). For example, drone technology with multispectral imaging sensors has recently been applied to characterise fungal disease in lemon myrtle trees (Heim et al., 2019) and macroalgal community structure in intertidal habitats (Tait et al., 2019), when combined with classification algorithms to disentangle biological communities. These recent findings suggest that drone technology may be applicable in many ecosystems, when combined with ground-truthed biodiversity data. However, biodiversity metrics are not the only ecological indicators, and functional traits are also widely used as ecological indicators of environmental change (e.g., Bjorkman et al., 2018). Because spectral reflectance is related to functional form (Gamon et al., 1997), the applicability of sensing technology is not limited to biodiversity, and can also act as a proxy for functional diversity (Frye et al., 2021). In the current study site, incorporating UAV monitoring to long-term experimental manipulations will enable the investigation of how environmental disturbances such as drought and nutrient addition influence biodiversity, functional diversity and community structure.

Another advantage of the current study is our use of raw digital radiance values and moments of spectral radiance distributions as opposed to ground-calibrated reflectance values that are typically applied to long-range multispectral imaging (Schläpfer et al., 2020). We therefore call for the application of the current approach across ecosystems. The public interest in drone technology may also assist in the development of crowd sourcing or citizen science programs to provide high-resolution biodiversity monitoring globally. Furthermore, the utility of this approach to gather large amounts of image data will increase when combined with machine learning algorithms to identify single species or high-resolution community dynamics. Currently, machine learning has been applied to agricultural imaging challenges (Heim et al., 2019) and in static species-cover assessments (Lopatin et al., 2017). Therefore, there is now an opportunity to combine this technology with large UAV-derived datasets. Ultimately, capitalising on drone technology has the potential to revolutionise biodiversity research.

Two key limitations to the current approach that may influence its application in other environments is the spatial and temporal resolution of data. Integrating spatial scales has long been a central issue in remote sensing applications (Turner, 2014). We found a negative association between spectral diversity and recording height. With increases in recording height there are decreases in pixel resolution, which are then unable to discern micro-structures from plant species that contribute to spectral reflectance variation. While this negative relationship was insufficient to influence species-diversity relationships in the current study below 10 m in recording height, this finding suggests that at greater recording heights associations with high-resolution measures of species diversity may breakdown. Previous studies have either focussed on short-range/destructive sampling (Frye et al., 2021; Lopatin et al., 2017) or long-range multispectral imaging (Gholizadeh et al., 2019), and therefore there is a need to explore species-spectral diversity associations across spatial scales. Recently, novel dissimilarity approaches have been applied to satellite imaging data at a range of spatial scales (Rossi et al., 2021). Integrating these approaches using UAV technology in novel habitats with alternate plant communities will be crucial for future research.

Temporal resolution is also a key factor in need of consideration when assessing spectral-species diversity associations. The data used in the current study represent a ‘static’ measure of biodiversity at the peak of the growing season in a temperate grassland. However, grassland communities exhibit a high degree of temporal variability, particularly in response to environmental drivers (Harrison et al., 2015; Thorhallsdottir, 1990). When implementing unmanned biodiversity monitoring, this temporal component may hinder the accuracy of spectral diversity measures. A solution developed by Rossi et al., (2021) applies a novel dissimilarity index between pairs of spectral images over the same region to disentangle temporal components of community change linked to management and phenology. However, this approach has yet to be applied to images collected using drone technology in varying habitats.

## 5. Conclusions

Taking advantage of technological advancements in unmanned sensing will greatly improve the cost-effectiveness of biodiversity monitoring. UAVs have the potential to span spatial scales, access challenging environments, and provide high resolution data on the impact of environmental change on ecosystems. Our study adds to a growing body of literature highlighting links between spectral and species diversity. Integrating these patterns at varying spatio-temporal scales and in novel habitats will provide vital insights to aid in documenting changes in the biosphere.

## Supporting information

Electronic Supplementary Information

## Data Availability Statement

We intend to submit all data to the dryad repository on acceptance, and archive code used in analyses using Zenodo.

## Credit author statement

The study was designed by JJ, RS-G, AH and NH. Core DroughtNet biodiversity data were collected and experimentation designed by CSL, AH, KM. DRAGNet biodiversity data and image data were collected by JJ, CA, EH, RH, LW, PF, and RS-G, with support from CSL. JJ analysed the data and wrote the manuscript, with initial feedback from RS-G. Further manuscript feedback was provided by CSL, KM, CA, LW and AH. All authors approved the manuscript for publication.

## Acknowledgements

We thank the many researchers making use of the RainDrop Wytham Woods site in summer 2021, who contributed vital discussions on botany and field logistics. We thank the Environmental Change Network, and in particular S. Schäfer and D. Pallett from the Centre for Ecology & Hydrology for access to the local weather data. Thanks to S. Middleton and D. Encarnation for support in botanical surveys. Special thanks to N. Fisher, N. Havercroft, and K. Crawford for field logistic support. JJ was funded by the Amazon Web Service Test Bed Funding scheme “Monitoring and Predicting Biodiversity Resilience through AI & Robotics” to RS-G and NH. RS-G was funded by a NERC IRF NE/M018458/1. NH was funded by EPSRC Programme Grant “From Sensing to Collaboration” (EP/V000748/1). AH was supported by the John Fell Fund. AH, KM and the Raindrop project were supported by the John Fell Fund, the Ecological Continuity Trust, the Patsy Wood Trust and the British Ecological Society.

