## Supplementary material for "Flexible estimation of biodiversity with short-range multispectral imaging in a temperate grassland": Electronic Supplementary Information

Supplementary Information Contents:


| | | |
|---|---|---|
| Fig. S1. | Site map of the Wytham Woods field site | 2 |
| Fig. S2. | Comparison of biodiversity metrics in ambient control treatments between DRAGNet and DroughtNet coordinated research network quadrats | 3 |
| Fig. S3. | Spectral coefficient of variation (and standard deviation) is a key indicator of biodiversity indices | 4 |
| Fig. S4. | Reduced spectral coefficient of variation in the drought and procedural control treatments | 5 |
| Fig. S5. | Structural interference on spectral reflectance of the examined grassland communities due to rainout shelters | 6 |
| Fig. S6. | Spectral coefficient of variation was consistent and repeatable at the quadrat-level | 7 |
| Table S1. | Model selection – Shannon-Weiner | 8 |
| Table S2. | Model selection- Simpson’s | 9 |
| Table S3. | Model selection – Band-level Shannon-Weiner models | 10 |
| Table S4. | Model selection – Band-level Simpson’s models | 11 |
| Table S5. | Model selection – Skewness and Biomass | 12 |
| References | | 12 |

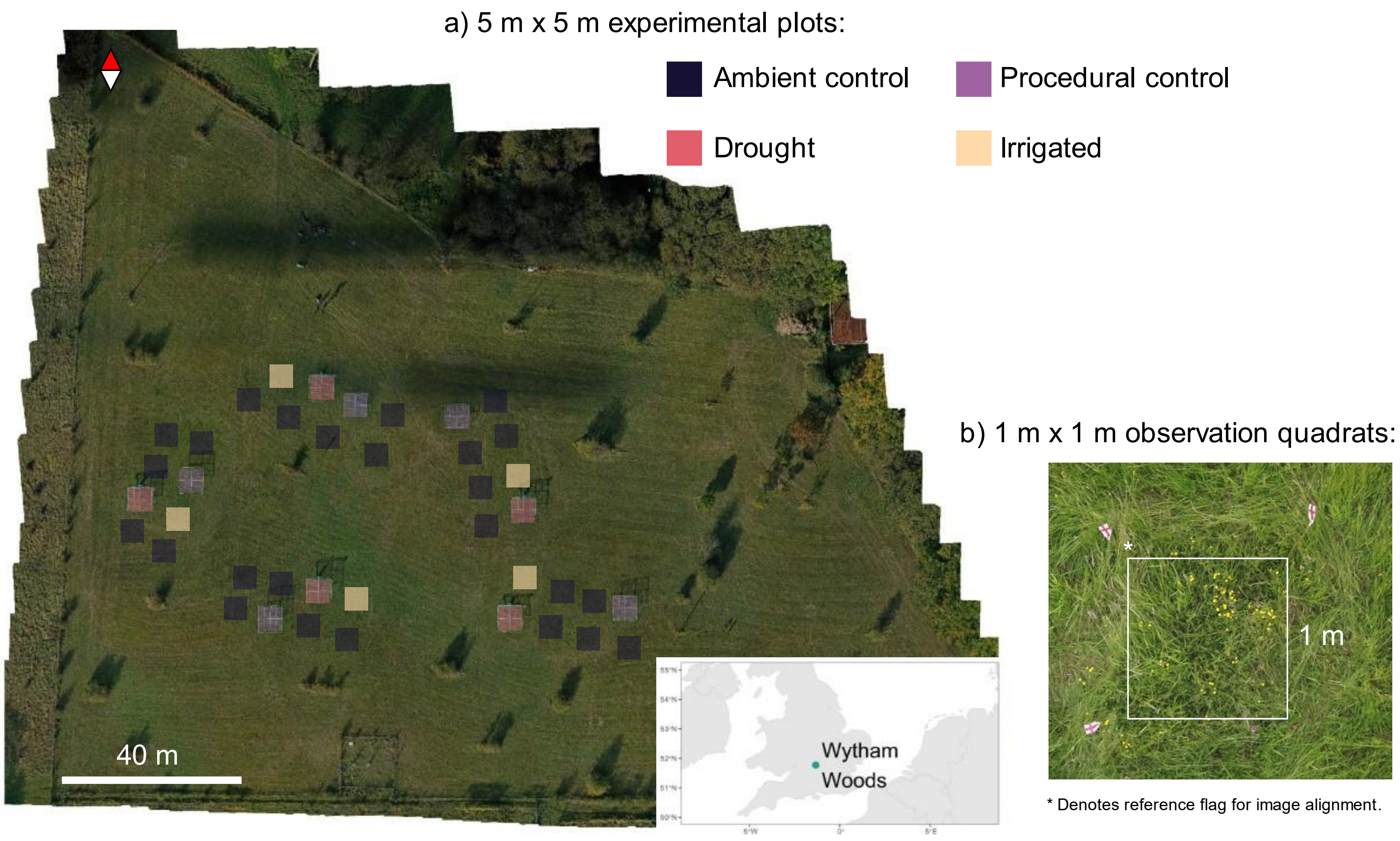


**Figure S1. Site map of the Wytham Woods field site with DRAGNet and DroughtNet coordinated research network plots (a) and observation quadrats for use in the study (b)**. a) Treatments include: ambient control plots (Ambient, n = 25), -50% rainfall shelters to simulate drought (Drought, n = 5), +50% irrigated plots to simulate increased rainfall (Irrigated, n = 5), and procedural controls (rainfall shelter with no change to rainfall) (Control, n = 2, 3 plots inaccessible for the UAV). b) example observation quadrat from the DRAGNet network. Flags were used to denote quadrat corners for UAV images, with an example denoted by the *.

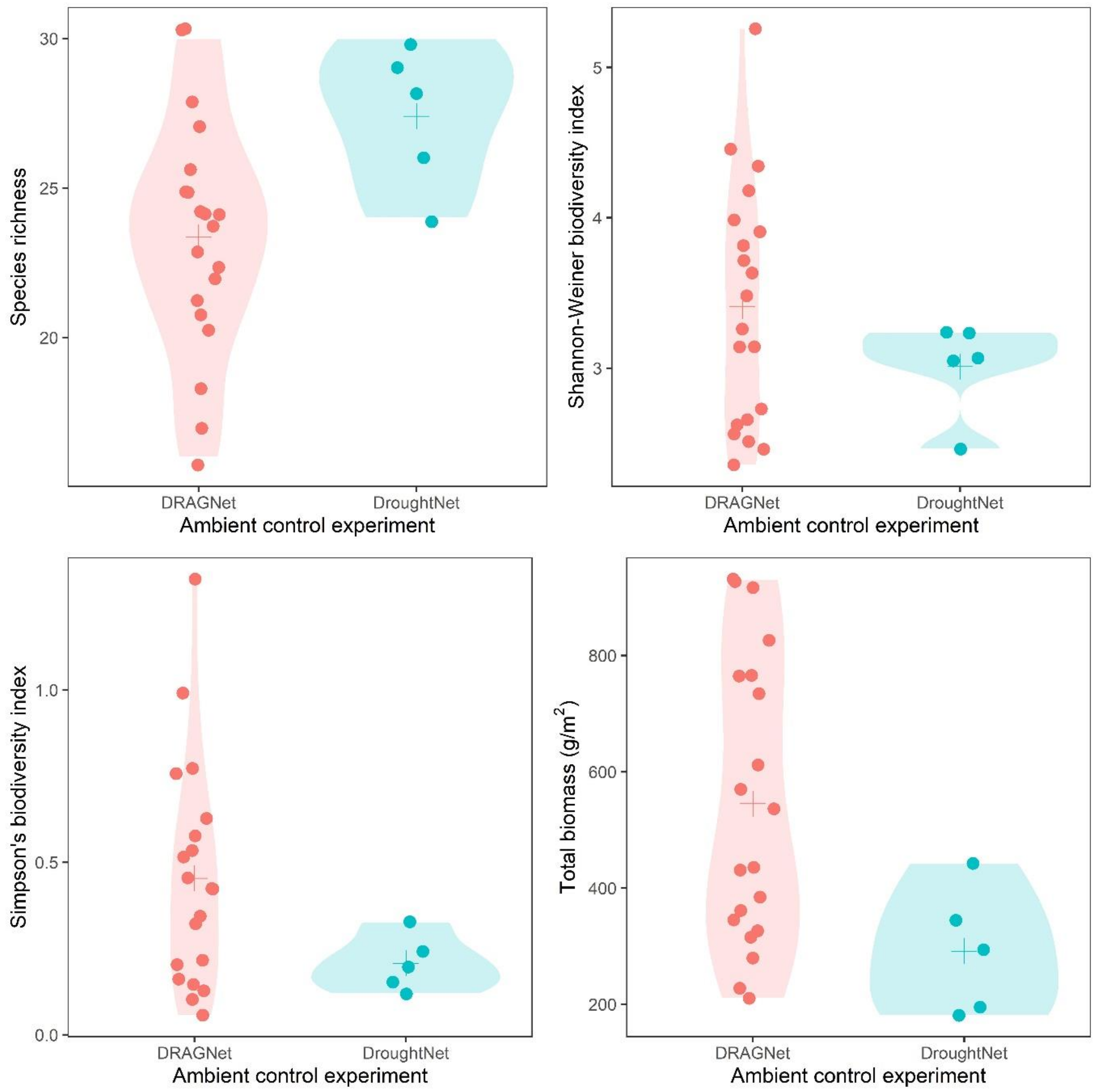


**Figure S2. Comparison of biodiversity metrics in ambient control treatments between DRAGNet and DroughtNet coordinated research network quadrats**. Each panel gives one of four biodiversity indices calculated from observation quadrats with respect to the coordinated research network, for ambient control quadrats only. Points are from observed quadrats for each group with its respective density summary. The + gives the mean value. We did not observe substantial significant differences between coordinated research networks in biodiversity indices (Shannon: $t = 1.75$, $df = 17.46$, $p = 0.10$; ln Simpsons: $t = 2.23$, $df = 14.1$, $p = 0.042$), but there was increased richness in DroughtNet Ambient control plots (Richness: $t = -2.93$, $df = 10.0$, $p = 0.02$) and decreased biomass(ln Biomass: $t = 2.88$, $df = 7.63$, $p = 0.02$). Nevertheless, as we did not observe strong differences between research networks, ambient control treatments were pooled in subsequent analyses.

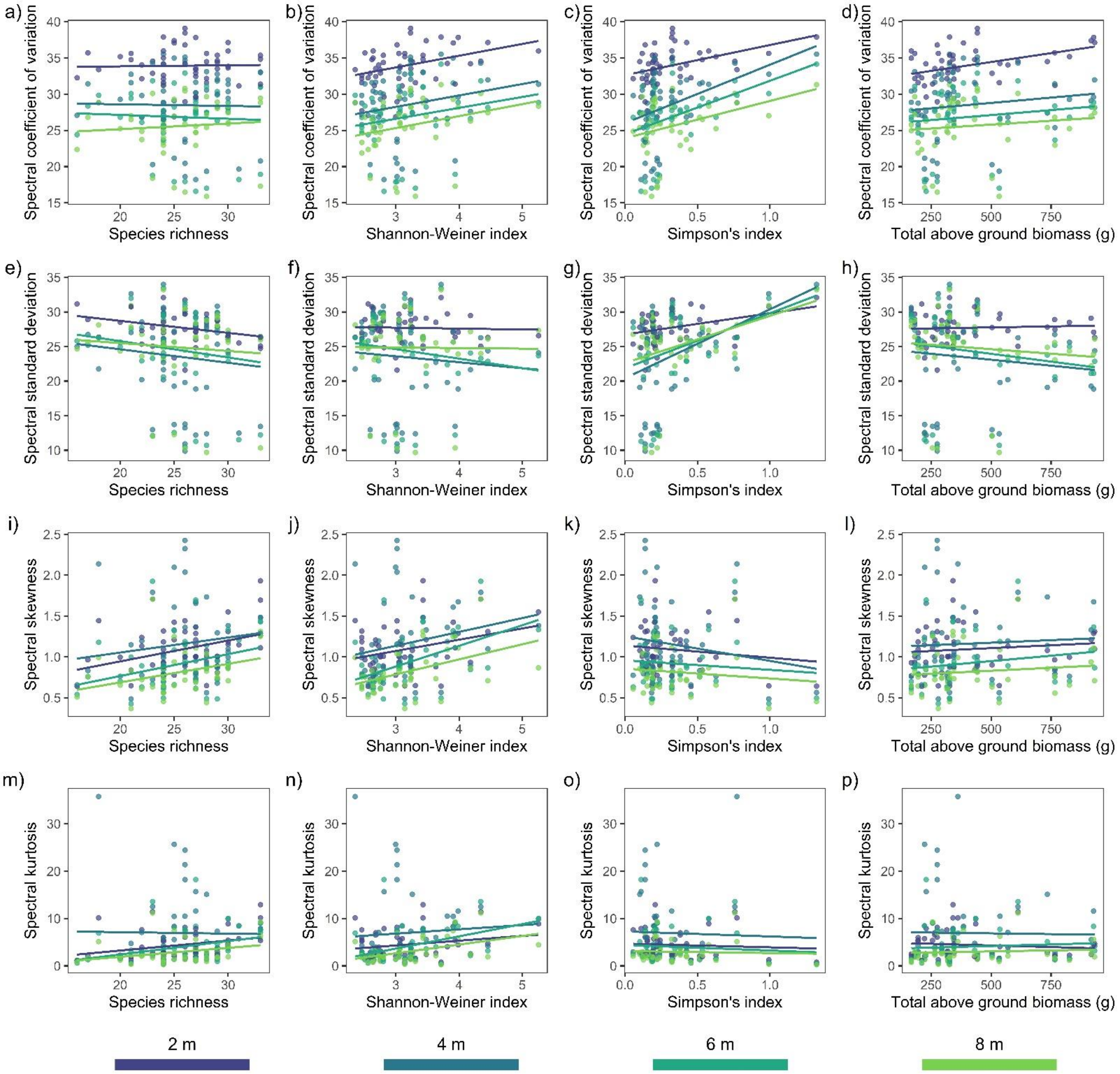


**Figure S3. Spectral coefficient of variation (and standard deviation) is a key indicator of biodiversity indices. Scatter plots of four biodiversity indicators with four multispectral moments**. Each panel provides a biodiversity indicator-multispectral moment value combination, where each observation is a plot at a given image recording height in a given sampling event. The colour of the point/line indicates the image recording height, for which there was a reduction in spectral moments with increased height. Lines are indicative linear regressions only to highlight general trends and were not estimated using Bayesian regression models.

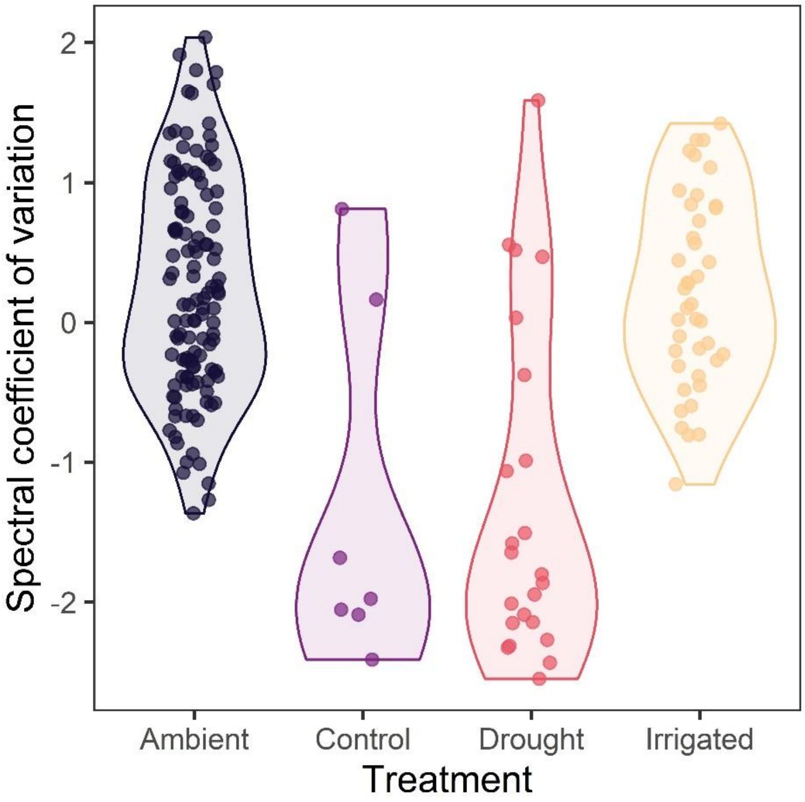


**Figure S4. Reduced spectral coefficient of variation in the drought and procedural control treatments.** Comparisons of spectral coefficient of variation between experimental treatment plots. Points correspond to field observations from a single plot at a different image recording heights, and the colour denotes the treatments from the Drought-Net and DRAGNet coordinated networks: ambient control plots (Ambient, n = 25), -50% rainfall shelters to simulate drought (Drought, n = 5), +50% irrigated plots to simulate increased rainfall (Irrigated, n = 5), and procedural controls (rainfall shelter with no change to rainfall) (Control, n = 2, 3 plots inaccessible for the UAV). All DRAGNet plots are classified as Ambient treatments, as the measurements were done prior to the application of any treatment – mechanical disturbance took place in Aug 2021, and fertilisation will take place in May 2022. Violin shapes indicate data density across spectral coefficient of variation values for each treatment. We observed reductions in spectral coefficient of variation for Control ($\beta_{\mathrm{TREATMENT,C}} = -1.39\,[-2.02, -0.72]$) and Drought ($\beta_{\mathrm{TREATMENT,D}} = -1.48\,[-1.90, -1.06]$) treatments, with predictive support relative to the base model ($\Delta\mathrm{elpd} = 9.44$).

a) b) c)

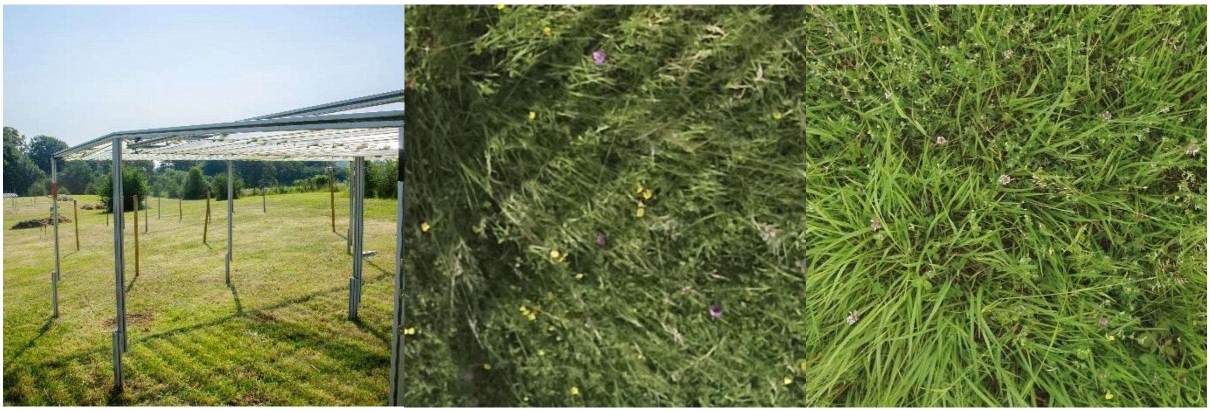

**Figure S5. Structural interference on spectral reflectance of the examined grassland communities due to rainout shelters.** These images are observations from the DroughtNet treatment plots. A key characteristic of the DroughtNet protocol is the use of metal rainfall shelters (a) to manipulate rainfall (Drought and Procedural control treatments). The RGB observation (2 m image recording height) from the Drought treatment (b) clearly displays visual interference in the form of shadows, which is caused by sunlight passing through the plastic rain gutters (seen in a). In contrast, the ambient control treatment image (c; 2 m image recording height) from the same observation day does not display this interference. Thus, reductions in spectral coefficient of variation in these treatments is caused by these structural interferences, *i.e.* from dark sections in the images that lower the variance in spectral reflectance.

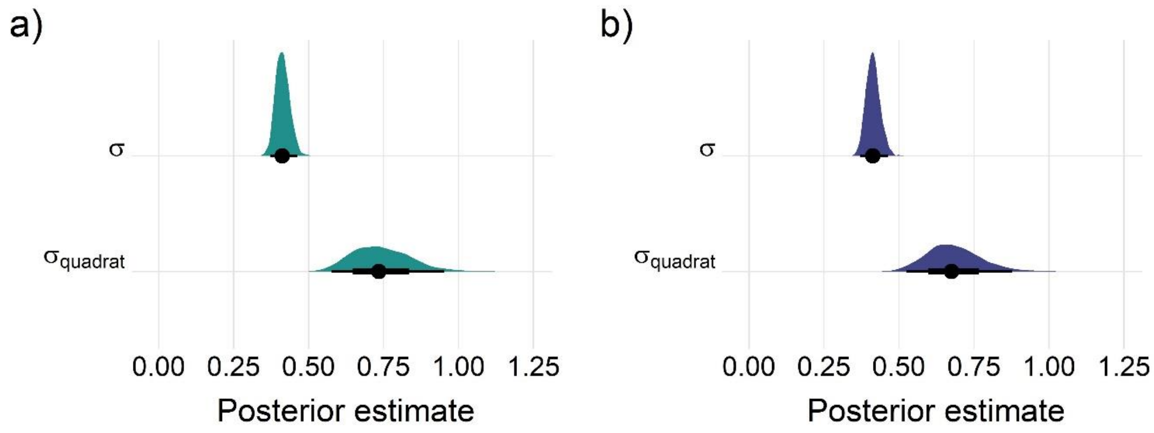


**Figure S6. Spectral coefficient of variation was consistent and repeatable at the quadrat-level across observation days and image recording height.** Posterior estimates of the quadrat-level random effect variance term $\sigma_{\text{quadrat}}$ with respect to the population-level variance σ for both (a) Shannon-Weiner and (b) Simpson's index models. In both cases, the $\sigma_{\text{quadrat}}$ greatly exceeded the population-level variance σ.

**Table S1. Increased model predictive performance including the Shannon-Weiner biodiversity index.** Model selection results for the relationship between coefficient of variation in spectral reflectance and the Shannon-Weiner biodiversity index. All models included an intercept-only random effect of the quadrat in addition to the predictor terms displayed. Model selection was performed with leave-one-out (LOO) cross validation and the expected logwise predictive density (elpd) from the loo R package (Bürkner, 2017). Most appropriate predictive model highlighted in bold. The term 'height' refers to image recording height.

| Model predictors | LOO elpd | LOO elpd error | $\Delta elpd$ | elpd error difference | LOO information criterion |
|---|---|---|---|---|---|
| height | -126.70 | 17.99 | 0.00 | 0.00 | 253.41 |
| shannon + height | -126.79 | 18.18 | -0.09 | 0.53 | 253.59 |
| **shannon + height + shannon:height** | **-127.70** | **17.91** | **-0.99** | **0.69** | **255.39** |
| shannon | -251.00 | 9.26 | -124.30 | 13.43 | 502.00 |
| base model | -251.26 | 9.31 | -124.56 | 13.47 | 502.53 |

**Table S2. Increased model predictive performance including Simpson's biodiversity index.** Model selection results for the relationship between coefficient of variation in spectral reflectance and Simpson's biodiversity index. All models included an intercept-only random effect of the quadrat in addition to the predictor terms displayed. Model selection was performed with leave-one-out (LOO) cross validation and the expected logwise predictive density (elpd) from the loo R package (Bürkner, 2017). Most appropriate predictive model highlighted in bold. The term 'height' refers to image recording height.

| Model predictors | LOO elpd | LOO elpd error | $\Delta elpd$ | elpd error difference | LOO information criterion |
|---|---|---|---|---|---|
| **simpsons + height + simpsons:height** | **-126.30** | **17.99** | **0.00** | **0.00** | **252.60** |
| simpsons + height | -126.60 | 18.13 | -0.30 | 1.03 | 253.19 |
| height | -126.70 | 17.99 | -0.41 | 1.22 | 253.41 |
| simpsons | -249.67 | 9.30 | -123.38 | 13.64 | 499.35 |
| base model | -251.26 | 9.31 | -124.96 | 13.52 | 502.53 |

**Table S3. Evidence for differences in the association of spectral-reflectance and the Shannon-Weiner biodiversity index between reflectance bands.** Model Selection results for the relationship between coefficient of variation in spectral reflectance and the Shannon-Weiner biodiversity index, separated across multispectral bands. All models included an intercept-only random effect of the quadrat in addition to the predictor terms displayed. Model selection was performed with leave-one-out (LOO) cross validation and the expected logwise predictive density (elpd) from the loo R package (Bürkner, 2017). Most appropriate predictive model highlighted in bold. The term 'height' refers to image recording height.

| Model predictors | LOO elpd | LOO elpd error | $\Delta elpd$ | elpd error difference | LOO information criterion |
|---|---|---|---|---|---|
| **shannon + band + shannon:band + height** | **-548.75** | **31.10** | **0.00** | **0.00** | **1,097.49** |
| shannon + height + band | -552.43 | 30.65 | -3.68 | 3.50 | 1,104.85 |
| shannon + height | -1,239.34 | 18.77 | -690.60 | 29.47 | 2,478.68 |
| base model | -1,320.73 | 19.91 | -771.98 | 30.48 | 2,641.46 |

**Table S4. Evidence for differences in the association of spectral-reflectance and Simpson's biodiversity index between reflectance bands.** Model Selection results for the relationship between coefficient of variation in spectral reflectance and Simpson's biodiversity index, separated across multispectral bands. All models included an intercept-only random effect of the quadrat in addition to the predictor terms displayed. Model selection was performed with leave-one-out (LOO) cross validation and the expected logwise predictive density (elpd) from the loo R package (Bürkner, 2017). Most appropriate predictive model highlighted in bold. The term 'height' refers to image recording height.

| Model predictors | LOO elpd | LOO elpd error | $\Delta elpd$ | elpd error difference | LOO information criterion |
|---|---|---|---|---|---|
| **simpsons + band + simpsons:band + height** | **-549.26** | **30.62** | **0.00** | **0.00** | **1,098.51** |
| simpsons + height + band | -552.92 | 30.70 | -3.67 | 2.13 | 1,105.85 |
| simpsons + height | -1,239.19 | 18.82 | -689.93 | 28.64 | 2,478.38 |
| base model | -1,320.73 | 19.91 | -771.47 | 29.64 | 2,641.46 |

**Table S5. No clear evidence for a relationship between above-ground biomass and the skewness in spectral reflectance.** Model Selection results for the relationship between the skewness of spectral reflectance and the above-ground biomass biodiversity index. All models included an intercept-only random effect of the quadrat in addition to the predictor terms displayed. Model selection was performed with leave-one-out (LOO) cross validation and the expected logwise predictive density (elpd) from the loo R package (Bürkner, 2017). The term 'height' refers to image recording height.

| Model predictors | LOO elpd | LOO elpd error | $\Delta elpd$ | elpd error difference | LOO information criterion |
|---|---|---|---|---|---|
| height | -181.75 | 23.03 | 0.00 | 0.00 | 363.49 |
| **biomass + height** | -182.16 | 23.20 | -0.42 | 0.61 | 364.33 |
| biomass + height + biomass:height | -182.73 | 23.05 | -0.99 | 0.54 | 365.46 |
| base model | -206.06 | 19.01 | -24.31 | 6.74 | 412.11 |
| biomass | -206.29 | 19.14 | -24.54 | 6.65 | 412.57 |
